# Bio-mimicked Leaf-Imprinted Topographies: Pattern Characterization and Cell Response

**DOI:** 10.64898/2026.08.03.742635

**Authors:** Dhruv Salot, Shital Yadav, Abhijit Majumder

## Abstract

Proper alignment of cells is crucial for functioning of various tissues such as skeletal muscle tissues, neural cells, adipose-derived stem cells, etc. Current *in-vitro* fabrication methods to replicate the cellular environment, e.g., photolithography and 3D printing, are not cost-effective and cannot capture the complexity of the surfaces to which these cells are exposed to. In this work, we used bio-mimicked leaf templates to closely resemble the *in-vivo* environment the muscle cells and cultured C2C12 cells, myoblast cell lines, on modified PDMS substrates fabricated using these leaf templates. Using image analysis software, we analyzed the degree of alignment of cells, aspect ratio and the area projected by individual cells cultured on these surfaces. The C2C12 cells cultured on the PDMS substrates formed utilizing the front and back sides of the leaves of *Musaceae Banana* were found to have an Aspect Ratio of 6.3 and 8.3, the highest among the surfaces studied in this paper. C2C12 cells cultured had the highest degree of alignment on the negative replica of the back side of Dracaena Sanderiana. Due to the availability of a wide range of leaf templates and bio-mimicked surface structures to measure cell response, it is difficult to find the optimal design. Hence, we have also tried to create a catalog using 15 different leaf surfaces and characterized these surfaces into various categories based on the grooves on the surfaces to provide a more comprehensive set of surface designs for studying cell behavior. To quantitatively analyze the groove pattern, 2D FFT analysis was also performed to find the dominant wavelength of the grooves. In surface characterization, hydrophobicity is also a parameter that needs to be considered; hence, the water contact angle of these surfaces was also measured. Our findings highlight the importance of surface topography and hydrophobicity in influencing cell alignment and can contribute to developing biomimetic surfaces for tissue engineering applications.

## 1. Introduction

The extracellular matrix (ECM) is a complex 3D network of macromolecules that provides structural support to cells along with giving necessary biochemical and mechanical cues essential for cell adhesion, migration, proliferation, differentiation, filtration and blood clotting (Frantz et al., 2010; Teti, 1992). ECM contains proteins such as collagen, laminin, and fibronectin that form diverse topographical features which include pores, ridges, fibers and other nanometer-scale features. Through self-assembly, these proteins organize into hierarchical structures spanning multiple length scales. For example, the ECM of the small intestine contains hierarchical structures ranging from the centimeter scale to the nanometer scale (Wang & L., 2011). The human skin dermis ECM under scanning electron microscope (SEM) reveals millimeter-sized ridges which are comprised of micrometer-scaled projections called dermal papillae, covered by sub-micron folds and pores (Kawabe et al., 1985). Another example is the hierarchical structures present on bones, ranging from submillimeter concentric cylinders called osteons to micrometer-scaled collagen fibers (Stevens & George, 2005). Previous studies have demonstrated that ECM topographies affect cellular behaviours such as cell adhesion (Biela et al., 2009), proliferation (Anselme & Bigerelle, 2011; Cai et al., 2006), migration (Diehl et al., 2005), differentiation (Dalby et al., 2006; Yang et al., 2014; Yim et al., 2007), morphology and orientation (Kim et al., 2009). To analyze the effect of ECM architecture on cellular functions, various 2D and 3D hierarchical structures have been prepared using microfabrication techniques such as photolithography (Molnar et al., 2007; Ross et al., 2012), capillary force lithography (Kim et al., 2009; Suh et al., 2009), electrospinning (Li et al., 2021; Su et al., 2021), electron beam lithography (Hatakeyama et al., 2007; Meyle et al., 1995), microcontact printing (Khadpekar et al., 2019; Wong et al., 2017) and 3D printing (Derakhshanfar et al., 2018). These methods, being highly precise, help fundamentally understand the effect of simple topographical features on cellular functions. For example, Molnar et al., 2007 investigated the influence of channel width on the differentiation of C2C12 cells into myotubes. However, such approaches are inherently limited in their ability to fabricate complex, multiscale topographical features of the ECM as commonly seen in-vivo. Although methods such as ion beam lithography, reactive ion etching (López-Bosque et al., 2013) and stereolithography 3D printing (Totonelli et al., 2012) have been utilized to develop complex hierarchical structures, their widespread adoption is limited by high cost and technical complexity.

In contrast to engineered microfabricated substrates, naturally occurring surfaces provide more topographical diversity and a large area for cell culture (Schulte et al., 2009). These surfaces have been utilized in earlier studies for various biomedical applications such as rose petal templates to capture circulating tumor cells (Dou et al., 2017), leaf templates for differentiation of neural stem cells (Hsu et al., 2017), cellular alignment of C2C12 cells (Yadav & Majumder, 2022), adhesion of cervical cancer cells (Zhang et al., 2022) and differentiation of adipose-derived stem cells (Ramaswamy et al., 2021). In these studies, soft lithography has been employed to bio-mimic the hierarchical micro- and nanoscale structures found on various plant surfaces, particularly leaves. Since leaf surfaces inherently exhibit complex topography, characterized by prominent primary veins at the macroscale, interconnected secondary vein networks at the microscale, and tertiary features such as trichomes (hairs) or waxy patterns spanning the micro- to nanoscale. Consequently, biomimicry of leaves results in surfaces with hierarchical features closely resembling those observed in-vivo tissues. For example, the superhydrophobic surface of lotus leaf had structures analogous to villi structures present in rat intestine (Quéré, 2008).

Since there are numerous natural templates, it is essential to investigate them further and create a repository that the researchers can use in the future as a reference for selecting surface patterns concerning objectives i.e., (i) a basic understanding of cellular response to unique pattern, (ii) controlling cellular behavior, (iii) use pattern with features similar to *in-vivo* tissue to mimic cellular microenvironment better. Vermeulen et al., 2021 proposed the potential application of topography on cellular behaviour by investigating 26 plant and insect surfaces.

In this work, we investigated the morphology of various leaf surfaces found locally. We observed different morphologies varying from simple to superimposed patterns, which are primarily complex in fabrication from currently available methods. Using soft lithography, we imprinted and characterized the surface morphologies obtained from leaf/plant surfaces on polymer polydimethylsiloxane, PDMS. The first imprint of the leaf surface on PDMS is termed a negative replica. The topological analysis of the fabricated negative replica on PDMS was done using a microscope and ImageJ software. We further investigated the wettability properties of fabricated patterns via biomimicking natural templates. Therefore, such exploration can give us unique patterns that mimic in-vivo morphology or help us understand cellular responses to complex patterns.

In vivo, geometrical cues provided by the ECM are critical regulators of cell alignment (Frantz et al., 2010). Unidirectional cellular arrangement is a defining feature for several tissues, including muscle, tendons and bones (Zheng et al., 2020). Previous studies have demonstrated that the differentiation of C2C12 myoblasts into myotubes is strongly influenced by simple topographical features of the underlying substrate including channels and tori (Bajaj et al., 2011). Accordingly, in this study, C2C12 cells were cultured on PDMS substrates with hierarchical structures to investigate the influence of multiscale topography on cellular alignment. Quantitative assessment of cellular alignment was performed using ImageJ. Cell morphology and orientation were characterized by measuring the aspect ratio, projected cell area, and orientation angle of individual cells relative to the underlying substrate topography. These metrics were used to evaluate the extent of contact-guidance-induced alignment and to quantify the degree of cellular orientation and morphological adaptation in response to the patterned surface features.

## 2. Material and methods

### 2.1. Preparation of patterned substrate

Leaves of 15 different plants were selected locally, with isotropic, anisotropic or reticulate arrangements of hierarchical structures (these arrangements have been explained later in sections 3.1 and 3.2). The natural surface, i.e., bio-template, was pasted on the petri dish using double-sided tape (figure 1.a). Polydimethylsiloxane (PDMS Sylgard 184 from Dow Corning), a widely used elastomer was mixed in ratios ranging from 10/1 (w/w) with a crosslinker and then degassed under vacuum. The uncured PDMS elastomer was poured on the template and allowed to solidify for 12 hours at 60 ^0^C. The solidified PDMS was peeled off (figure 1.b) and the negative replica of the pattern was obtained. To have a control, flat PDMS, i.e., without a pattern, was prepared by pouring uncured elastomer into a 60 mm diameter flat petri dish and following the aforementioned method. There are numerous leaves with reticulate and branched veins. However, most leaves possess a hairy surface. During the PDMS imprinting process, these surface hairs tend to fracture and become embedded within the PDMS (as shown by yellow arrows in figure 1.d-e). Consequently, leaves with hairy surfaces are excluded from cell culture studies.

**Figure 1:**
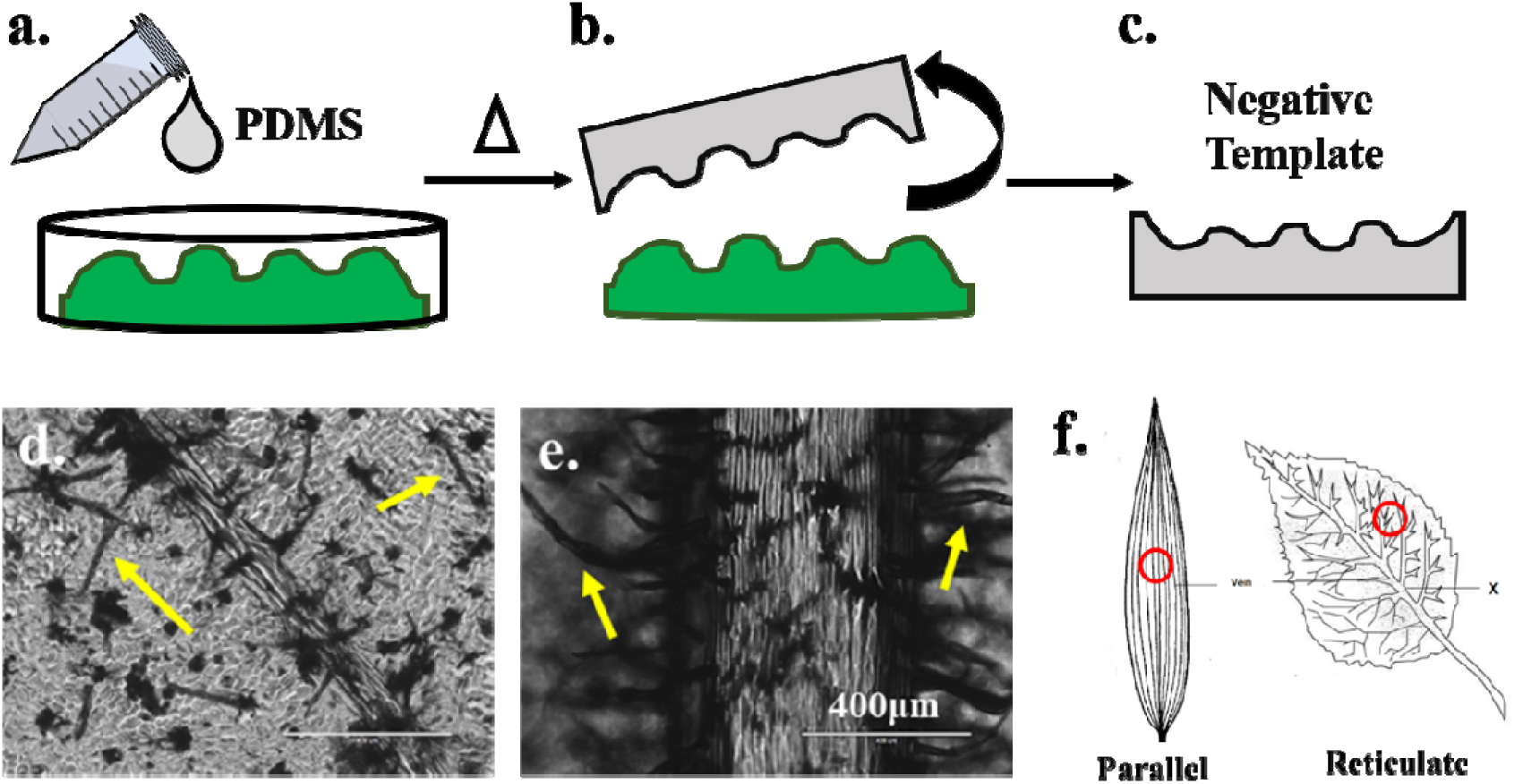
Schematic for steps involved in patterning PDMS from leaf pattern, (a-b) pouring PDMS on bio-template followed by crosslinking to obtain (c) negative replica; (d-e) images of hair-like structure obtained on PDMS pattern from hairy leaf and (f) classification of leaves used in this report i.e., parallel and reticulate venation leaves (red-area selected for imaging)

### 2.2. Fast Fourier Transform (FFT)

The characteristic wavelength (λ) of the anisotropic pattern was determined using the two-dimensional fast Fourier transform (2D-FFT) in FIJI software. The two-dimensional FFT function provides “frequency” space in the form of the rate of change of pixel intensity in the raw image (Bajaj et al., 2011). In the resultant FFT output, the closest high-frequency pixel point from the center was referred to as the dominant wavelength of the image. (Ayres et al., 2008)

For anisotropic continuous and discontinuous grooves there are 2 dominant wavelengths, parallel and perpendicular to the grooves (figure 2). These 2 wavelengths were found by simply drawing a line across the grooves in the above-stated directions using ImageJ and plotting the graph of pixel intensity vs location. The ratio of several peaks to the length of the line gives us the wavelength.

**Figure 2.**
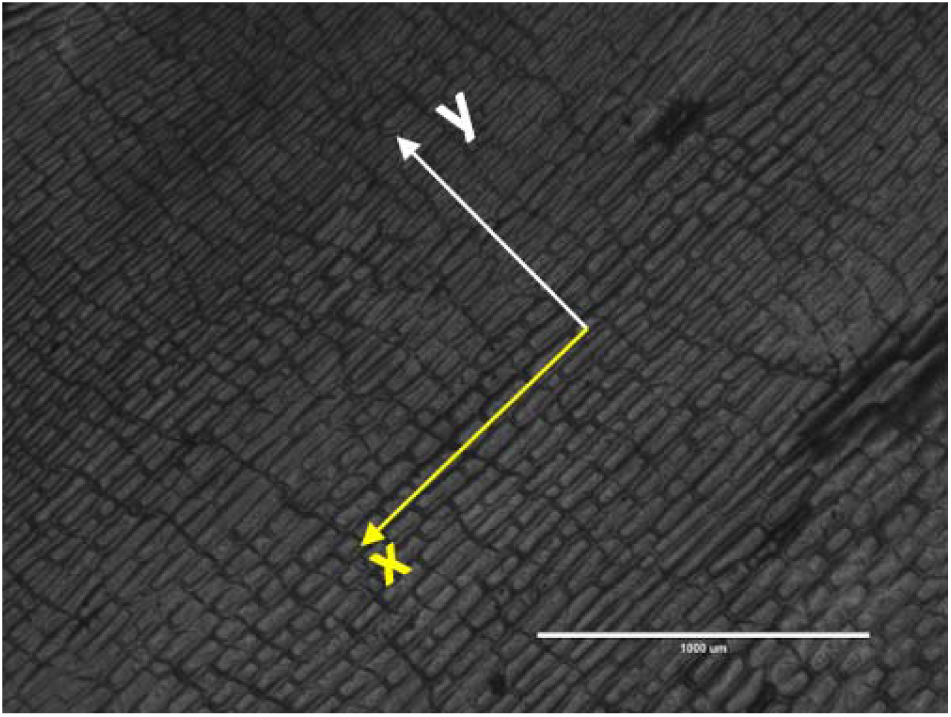
Image of the front side of leaf of *Yucca Gloriosa*. The x-axis in yellow represents the direction parallel to the grooves (indicated as “para” in subsequent images), and the y-axis in white represents the direction perpendicular to the grooves (indicated as “perp” in subsequent images). (Scale bar: 1000 m)

### 2.3. Water contact angle measurement

To measure the contact angle of water respective to prepared topographies, 5μl of DI water was pipetted on the substrate. The water droplet was captured from the side of the camera, and the contact angles were quantified using ImageJ software.

### 2.4. Cell Culture

To study the effect of topography on cells, we have cultured C2C12 (mouse myoblast skeletal cells) on the PDMS substrates. C2C12 cells were cultured in high glucose Dulbecco’s modified eagle medium (Himedia AL007A) supplemented with 1% Antibiotic-Antimycotic (Himedia-A002) solution, 1% L-glutamine (Gibco) and 20% fetal bovine serum (Himedia-RM1112). Cells were then trypsinized using a solution of trypsin-EDTA 1X (Himedia-TCL0033) and incubated for 5 minutes at 37°C. The suspension is centrifuged at 1000 rpm for 5 minutes to obtain a cell pellet. Cells were resuspended in a fresh medium and counted using a haemocytometer. The PDMS patterns were cut in a circular shape of 18mm diameter and placed in 12 well plates after which these substrates were sterilized with ethanol, followed by washing with Dulbecco’s phosphate buffered saline (DPBS Himedia-TS1006) to wash out the ethanol. These substrates were further treated with UV light for 30 minutes to sterilize the surface and were treated with plasma for 2 minutes, followed by rinsing thrice with Dulbecco’s phosphate-buffered saline (DPBS Himedia-TS1006). Collagen solution of 25 g/ml (diluted with PBS from a stock solution of 3mg/ml) was poured on the substrate. We placed it in an incubator overnight at 4°C to allow the collagen to spread over the surface uniformly. PDMS substrates were rinsed again with DPBS before C2C12 cells were then cultured on these surfaces using a cell density of 210-280 cells/cm² inside a laminar hood and placed in an incubator at 37°C. After 24 hours, the cells were stained with Calcein-AM, propidium iodide (PI), and Hoechst for imaging. These stained cells were imaged using ImageJ software (NIH) at 4x and 10x magnification.

### 2.5 Morphological analysis

Cells were stained with Calcein-AM (Invitrogen-C3099) and Hoechst (Invitrogen-H3570) after 24h and then imaged under an inverted microscope (EVOS-FL Auto, Life Technologies) at 10x magnification. The images were analysed to obtain the cell area and aspect ratio using the ImageJ software. The cellular projected area was determined by manually tracing the perimeter of a single cell. Tracing of the perimeter in ImageJ was also used to determine the aspect ratio of a single cell, calculated as follows:

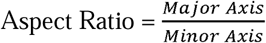

A total of 30 patterned PDMS substrates were fabricated, of which 10 substrates exhibiting an anisotropic groove pattern with a parallel vein-like structure were selected for morphological analysis. For each of these 10 substrates, measurements were performed on 60 single cells, and the average values of cell projected area and aspect ratio were reported.

### 2.6 Orientation Angle

Cellular orientation was measured as the angle of the entire cell orientation with respect to the groove direction in patterned PDMS and with respect to the x-axis in flat substrates using ImageJ. Cells are defined as oriented or aligned along the topography if the angle is less than 30°.

### 2.7 Statistical analysis

The data were plotted using GraphPad Prism. Differences were considered statistically significant at ∗p < 0.05, ∗∗p < 0.0001, ns = not significant using Student’s t-test or a one-way ANOVA.

### 2.8 Dominant Wavelength Estimation

The dominant wavelength of a pattern indicates the distance between consecutive grooves. For anisotropic, parallel-grooved patterns, the dominant wavelength captures groove spacing both parallel and perpendicular to the pattern. For the leaves in this study, the spacings range from approximately 50–400 µm parallel and 10–100 µm perpendicular to the grooves, which closely match the dimensions of C2C12 cells cultured on flat PDMS substrates (∼70–200 µm in length and 10–50 µm in width) (Yadav & Majumder, 2022). Consequently, the dominant wavelength serves as a meaningful descriptor for cell-scale guidance, as it indicates whether individual C2C12 cells can physically align with the grooves. Such alignment is critical for studying contact-guided cells.

## 3. Results

A wide variety of plant leaves were used as templates to imprint their surface features onto PDMS, resulting in a diverse repository of bio-inspired patterns, as described in the Materials and Methods section. Leaves are broadly categorized based on their venation into parallel-veined and reticulate-veined types (fig. 1.i), and this is also the classification used in this catalogue. Surface morphologies obtained on PDMS by negative replica from the front and back side of leaves were examined as discussed in the following sections.

### 3.1. Leaves with parallel veins

In this category of leaves, veins are aligned axially and are non-intersecting (figure 1.i).

#### 3.1.1 Isotropic pattern

In this work, leaves with uniform pattern with no significant directionality is considered as isotropic pattern. For example, patterns from the front and back side of *Codiaeum Variegatum* (fig 3.a-b) fall under this category for the following reasons stated. First, the grooves observed on these surfaces (fig 3.a.i-b.i) are irregular in shape and have no preferred orientation. Second, these grooves are distributed uniformly across the surface, resulting in an overall isotropic pattern. This is further confirmed by the FFT images (fig 3.a.ii-b.ii), which display concentric rings of high-intensity pixels originating from the central white spot indicating a direction-independent pattern. Since the surfaces in this type of pattern have no direction to their grooves, we have reported only one dominant wavelength for each surface (fig. 3.c). The dominant wavelength of (a) front and (b) back of *Codiaeum Variegatum* is 26 ± 2 m and 31 ± 6 m, respectively.

**Figure 3.**
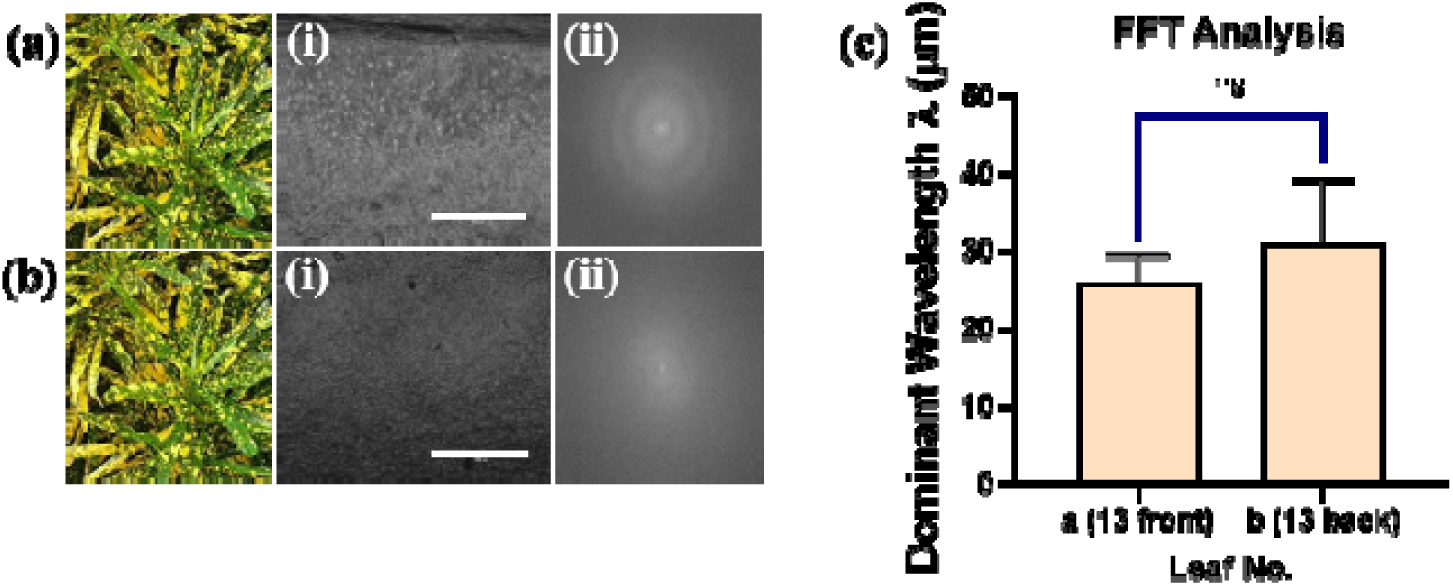
Images of (a) Leaf 13 front, (b) Leaf 13 back (refer to Table 1), (i) Microscope images of PDMS substrate (Scale bar = 1000 µm) and (ii) FFT of respective PDMS substrate; (c) dominant wavelength of patterns (m) (**p < 0.0001, ns: not significant)

**Table 1.** Legend of leaf surfaces.

| Sr.No. | Leaf name | Reference image |  |
| --- | --- | --- | --- |
|  |  | Front | Back |
| 1 | Plectranthus Scutellarioides | fig. 6.a.i | fig. 6.b.i |
| 2 | Musaceae Banana | fig. 4.e.i | fig. 5.e.i |
| 3 | Bauhinia Tomentosa | fig. 6.e.i | fig. 6.f.i |
| 4 | Yucca Gloriosa | fig. 4.f.i | fig. 5.f.i |
| 5 | Syngonium Podophyllum | fig. 6.g.i | fig. 6.h.i |
| 6 | Hibiscus Rosa-sinensis | fig. 6.k.i | fig. 6.l.i |
| 7 | Jasminum Polyanthum | fig. 6.i.i | fig. 6.j.i |
| 8 | Pseuderanthemum Carruthersii | fig. 6.m.i | fig. 6.n.i |
| 9 | Arecaceae | fig. 5.g.i | fig. 4.g.i |
| 10 | Dracaena Sanderiana | fig. 4.c.i | fig. 4.d.i |
| 11 | Cordyline Fruticosa | fig. 5.c.i | fig. 5.d.i |
| 12 | Tradescantia Zebrina | fig. 4.b.i | fig. 5.b.i |
| 13 | Codiaeum Variegatum | fig. 3.a.i | fig. 3.b.i |
| 14 | Tradescantia Spathacea | fig. 4.a.i | fig. 5.a.i |
| 15 | Erigeron Annuus | fig. 6.c.i | fig. 6.d.i |

#### 3.1.2 Anisotropic continuous pattern

Leaves in this category exhibit directional structures distributed uniformly over the surface, unlike isotropic pattern. Figure 4 summarizes leaves and respective surface morphologies obtained on PDMS, which resemble anisotropic continuous patterns; these include the front side of *Tradescantia Spathacea* (fig 4.a), the front side of *Tradescantia Zebrina* leaf (fig 4.b), the front and back side of *Dracaena sanderiana* (fig 4.c and 4.d), front side of *Musaceae Banana* (fig 4.e), front side of *Yucca Gloriosa* (fig 4.f) and back side of *Arecaceae* (fig 4.g). The reason for including specifically these patterns in this category is following. First, these surfaces consist of rectangular grooves repeated throughout the surface and at regular intervals. These patterns are commonly observed on the front surface of leaves, where the veins are less pronounced and more widely spaced, resulting in a lamina dominated by a single type of grooves and yielding a uniform, continuously grooved surface. Second, the FFT (fig. 4) of these surfaces usually consists of bright pixels arranged as a narrow band across the image which accounts for the directionality of the grooves and repeating white spots along these bands depict the repetition of the grooves throughout the surface.

**Figure 4.**
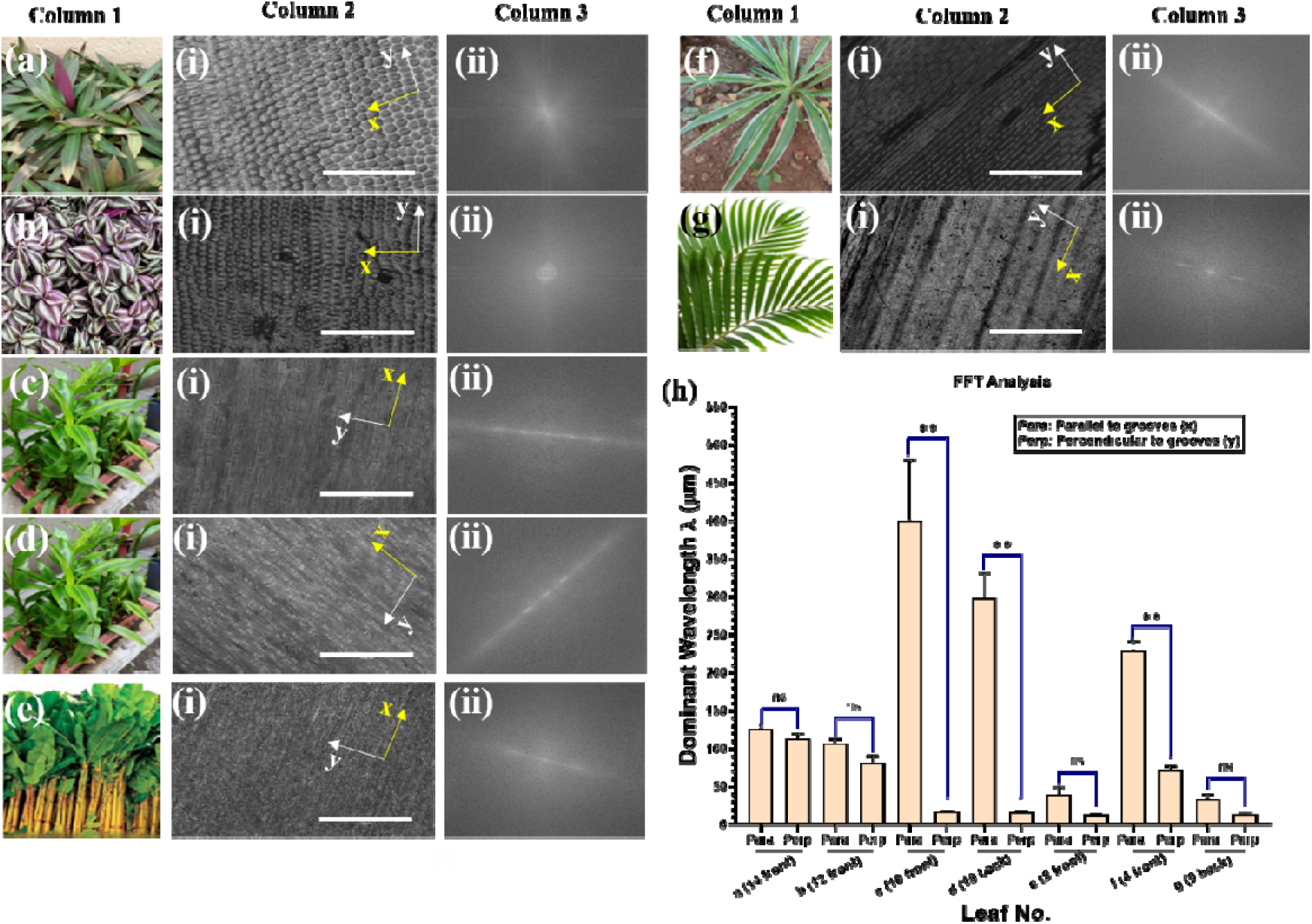
Column 1: leaf images of (a) Leaf 14 front, (b) Leaf 12 front, (c) Leaf 10 front, (d) Leaf 10 back, (e) Leaf 2 front, (f) Leaf 4 front, (g) Leaf 9 back with respective, Column 2: Microscope images of PDMS substrate (yellow x: parallel to grooves, white y: perpendicular to grooves; Image scale bar = 1000 µm) and Column 3: FFT of respective microscope images. (h) Dominant wavelength (λ) of leaves parallel and perpendicular to the grooves **p < 0.0001, ns = not significant.

The topography of *Tradescantia Spathacea* and *Tradescantia Zebrina* features square or hexagonal patterns arranged at regular intervals. The FFT analysis of these 2 surfaces (fig. 4.a.ii – 4.b.ii) displays a circular pattern, similar to the isotropic pattern described in the previous category. However, due to the repeating white spots in the FFT images (fig 4.a.ii – 4.b.ii), rather than continuous concentric rings, indicate that the spatial pattern is not uniformly distributed across all directions therefore, these surfaces cannot be classified in the previous isotropic category. Using the fig 4.a.i and 4.b.i, we find dominant wavelength in both directions, i.e. parallel and perpendicular to the grooves giving a value of 126 ± 6 µm and 113 ± 5µm respectively for *Tradescantia Spathacea* and 107 ± 4µm and 81 ± 7µm respectively for *Tradescantia Zebrina.* In the case of *Dracaena Sanderiana*, both the front (fig 4.c.i) and back (fig 4.d.i) surfaces exhibit hierarchical structures in the form of elongated rectangles. Among all the “anisotropic continuous patterns” analyzed, *Dracaena Sanderiana* has the highest dominant wavelength along the parallel direction (298 ± 24µm for the back side and 400 ± 67µm for the front). Front surfaces of *Musaceae Banana* (fig 4.e.i) and *Arecaceae* (fig 4.g.i) have the smallest rectangular grooves among those studied in this category. This is further supported by the graph in fig. 4.h, with the dominant wavelength along the parallel direction of each leaf surface being 40 ± 7µm and 33 ± 5µm respectively. Along with *Dracaena Sanderiana*, front side of *Yucca Gloriosa* (fig 4.f.i) has rectangular grooves with significant difference (>140µm) in the size of the grooves along the parallel and perpendicular direction (fig. 4.h). Dominant wavelength for the front side of *Yucca Gloriosa* (fig. 4.f.i) in the parallel direction is around 229 ± 11µm and along the perpendicular direction is 71 ± 7µm.

#### 3.1.3 Anisotropic discontinuous pattern

In this category, the patterns are intermittent, i.e. the regular arrangement of rectangular grooves in the lamina of the leaf is not uniform and has disruptions in the pattern, due to numerous stomata or intermittent veins. Figure 5 summarizes leaves and respective surface morphologies obtained on PDMS, which resemble anisotropic discontinuous patterns which include the back side of *Tradescantia Spathacea* (fig 5.a), the back side of *Tradescantia Zebrina* (fig 5.b), front and back side of *Cordyline fruticosa* (fig 5.c-d), back side of *Musaceae Banana* (fig 5.e), back side of *Yucca Gloriosa* (fig 5.f) and back side of *Arecaceae*. The rationale for this classification is as follows. First, the FFTs (fig.5 column 3) corresponding to this class of patterns predominantly exhibit a bright narrow band of pixels across the image, which is indicative for the directionality of the pattern. But unlike the previous category, these FFTs do not display repeating bright spots, providing evidence of discontinuities within the spatial pattern. Second, examining the corresponding microscope images (fig. 5 column 2) confirms the presence of rectangular groove features aligned along a preferred direction, thereby imparting anisotropic nature to the surface. However, the presence of intermittent veins that are more pronounced and more closely spaced than those in the previous category disrupts the regularity of the grooves, resulting in a non-uniform surface topography. Consequently, this type of pattern is typically observed on the back surface of leaves, where multiple stomata and more prominent intermittent veins contribute to the disruption in the groove pattern.

**Figure 5.**
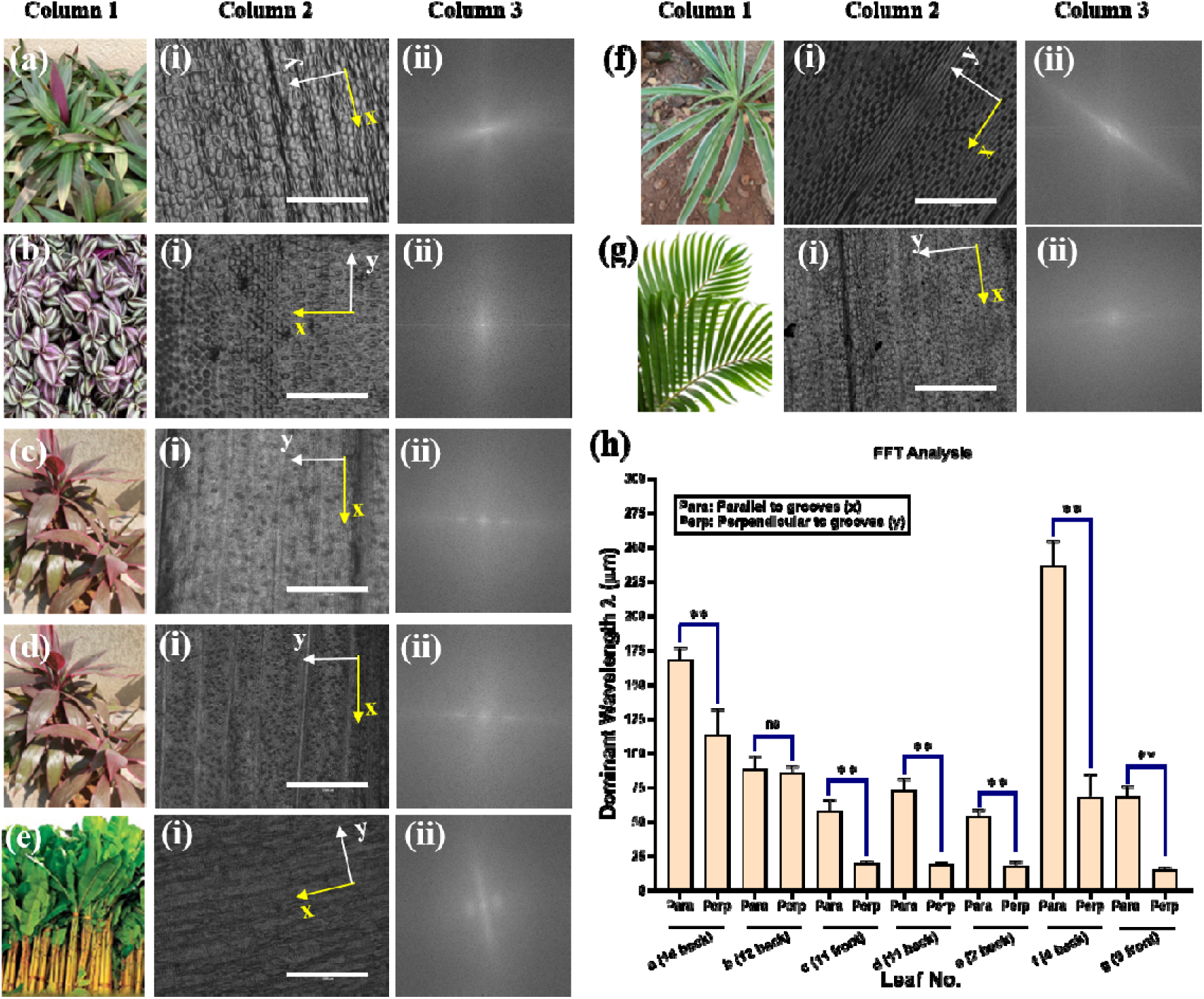
Column 1: Images of leaf of (a) Leaf 14 back, (b) Leaf 12 back, (c) Leaf 11 front, (d) Leaf 11 back, (e) Leaf 2 back (f) Leaf 4 back (g) leaf 9 front; Column 2: Microscope images of PDMS substrates of respective leaves (yellow x: parallel to grooves, white y: perpendicular to grooves; Image scale bar = 1000 µm); Column 3: FFT of respective images; (h) dominant wavelength of patterns (**p <0.0001, ns = not significant)

Within this category, the individual surfaces exhibit distinct topographical features. On the back side of *Tradescantia Spathacea* (fig 5.a.i) rectangular grooves are arranged irregularly as a result of the uneven topography of the leaf. This is further reflected in the leaf’s FFT (fig 5.a.ii), which exhibits a narrow band of high-intensity pixels, indicating directional alignment of the grooves, but the absence of repeating bright spots reveals a discontinuous pattern. Similarly, rectangular grooves are observed on the back side of *Tradescantia Zebrina* (fig 5.b.i), although the pattern is interrupted multiple times due to hairs and stomata. The grooves observed on the back side of *Tradescantia Zebrina* exhibit minimal variation between the parallel (88 ± 9µm) and perpendicular (85 ± 6µm) direction, resulting in more square-like grooves. As compared to the back side of *Tradescantia Spathacea*, having a significant difference of approximately 50µm. Front and back sides of *Cordyline fruticosa* leaves (fig 5.c.i-d.i) consist of similar rectangular grooves, the dominant wavelengths for the front side were 57 ± 10µm (parallel); 19 ± 3µm (perpendicular) and for the back side were 72 ± 9µm (parallel) and 18 ± 2µm (perpendicular). The FFT of both these surfaces (fig. 5.c.iii-d.iii) shows 3 consecutive white spots, but due to the uneven distribution of the stomata on the surface (more concentrated on the back side), these grooves are discontinuous, justifying their classification in this category. The back side of *Musaceae Banana* (fig. 5.e.i) has irregular rectangular grooves with a dominant wavelength of 54 ± 5µm in the parallel direction and 18 ± 4µm in the perpendicular direction, which is one of the lowest wavelengths among the leaves of this category studied. Back side of *Yucca Gloriosa* (fig. 5.f.i) consists of elliptical and rectangular shaped grooves at regular intervals, this is evident from The FFT of this leaf (fig 5.f.iii) the central white spot is surrounded by a bright ring depicting the uniformity of the pattern in the surface. But due to the veins on the surface the pattern is disrupted and in the FFT we do not have any repeating bright spot. Among the leaves studied in this category *Yucca Gloriosa* has the highest dominant wavelength i.e. 236 ± 20 m in the parallel direction and 68 ± 18 m in the perpendicular direction. For the front side of *Arecaceae* (fig 5.g.i) the surface is covered with irregular patterns and with rectangular grooves of the veins. In the parallel direction the dominant wavelength is 68 ± 7 m and in the perpendicular direction is 15 ± 2 m. Due to the veins of the surface and directionality of the pattern, the FFT contains a faint bright band even though the patterns are irregular.

### 3.2. Leaves with reticulate veins

Leaves with reticulate veins (fig 1.f) comprise two regions, (i) vein region and (ii) area between the veins, referred to as stomata region in this work. The stomata region showed different morphologies, whereas the vein region is nearly similar in all cases. Another characteristic of this leaf category is that the FFT analysis (figure 6, column 3) exhibits a uniform circular distribution of pixels around the origin, indicating that the groove patterns are neither pronounced nor periodically repeating. Consequently, the dominant wavelength for these patterns was not determined.

**Figure 6.**
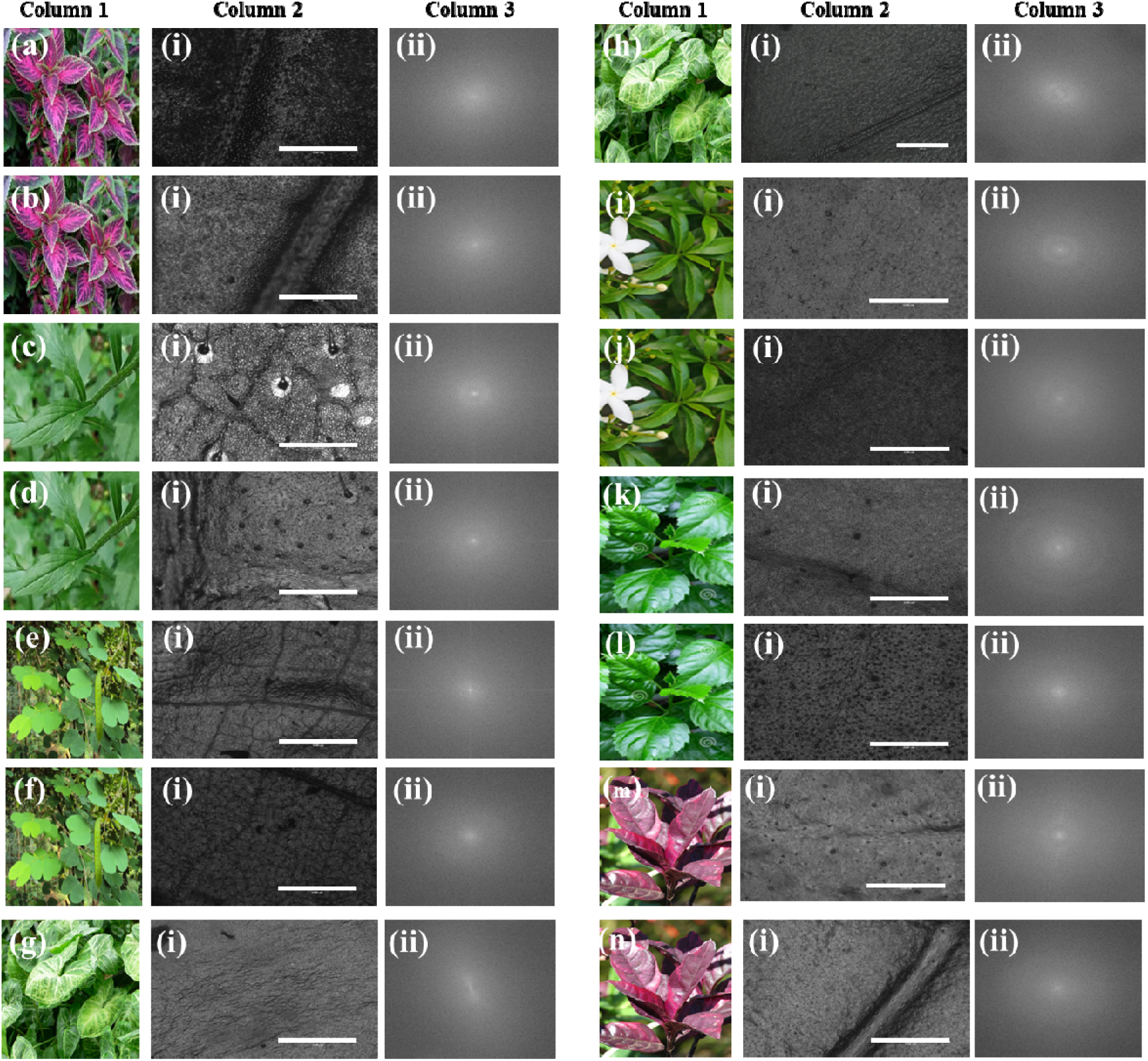
Column 1: Images of the leaf of (a) Leaf 1 front, (b) Leaf 1 back (c) Leaf 15 front, (d) Leaf 15 back, (e) Leaf 3 front, (f) Leaf 3 back, (g) leaf 5 front, (h) Leaf 5 back, (i) Leaf 7 front, (j)leaf 7 back, (k) Leaf 6 front, (l) Leaf 6 back, (m) Leaf 8 front, (n) Leaf 8 back; Column 2: microscope images of respective leaves and Column 3: FFT analysis (microscope image scale bar = 1000 µm)

Figure 6 summarizes morphologies obtained from the front and back sides of 6 reticulate vein leaves. The stomata region of the front and back side of *Plectranthus Scutellarioides* (fig 6.a.i-6.b.i) exhibit numerous circular grooves, while the vein region displays rectangular grooves (fig 6.a.i-6.b.i) with abundant hairs. In the case of the pattern from *Erigeron Annuus* leaf, the front and back sides (fig 6.c.i-6.d.i) exhibit a limited number of elliptical or grain-like surface features, accompanied by a dense distribution of trichomes and sparsely distributed stomata openings. The rectangular grooved pattern is present in the vein region in both cases (fig 6.c.i-6.d.i). The front and back sides of the *Bauhinia tomentosa* leaf exhibit a highly interconnected vein network (fig 6.e.i-f.i.), with the stomata region show no pronounced surface undulations and densely covered with hairs. Several templates displayed an almost smooth surface morphology, as observed on the front surfaces of *Syngonium podophyllum* (fig 6.g.i), *Jasminum Polyanthum* (fig 6.i.i) and *Hibiscus Rosa-sinensis* (fig 6.k.i). The back side of *Syngonium podophyllum* leaf (fig 6.h.i) circular groove-like features are observed across both stomatal and vein regions, back side of *Jasminum Polyanthum* (fig 6.j.i) and *Hibiscus Rosa-sinensis* (fig 6.l.i) have circular/elliptical grooves in the area between the veins due to the presence of stomata. Front (fig 6.m.i and 6.n.i) and back side of *Pseudernthemum Carruthersii* show almost no undulations in the area between veins and have line-like or rectangular grooves within the veins. The FFT analysis of all the above leaves has a uniform distribution of grayscale intensity of pixels, indicating no significant patterns present for the cells to detect when cultured.

### 3.3. Wettability analysis

The wettability of cured PDMS is known to get altered based on the pattern or morphology of its surface (Kameya, 2017). Hydrophobicity of the surface on which cells are cultured plays a crucial role in the cell-to-surface interaction affecting cellular properties of cell adhesion (Ferrari et al., 2019), protein absorption (Falde et al., 2016) and other biomedical applications for regenerative medicines. Therefore, we checked the variations in the water contact angle as a measure of the hydrophobicity of the PDMS substrate after bio-mimicking leaf templates. Figure 7 shows the various substrates with respective contact angles. The contact angles measured on nearly all replicated topographies were higher than that of flat PDMS (95°), with the exception of the front side of *Erigeron annuus*.

**Figure 7:**
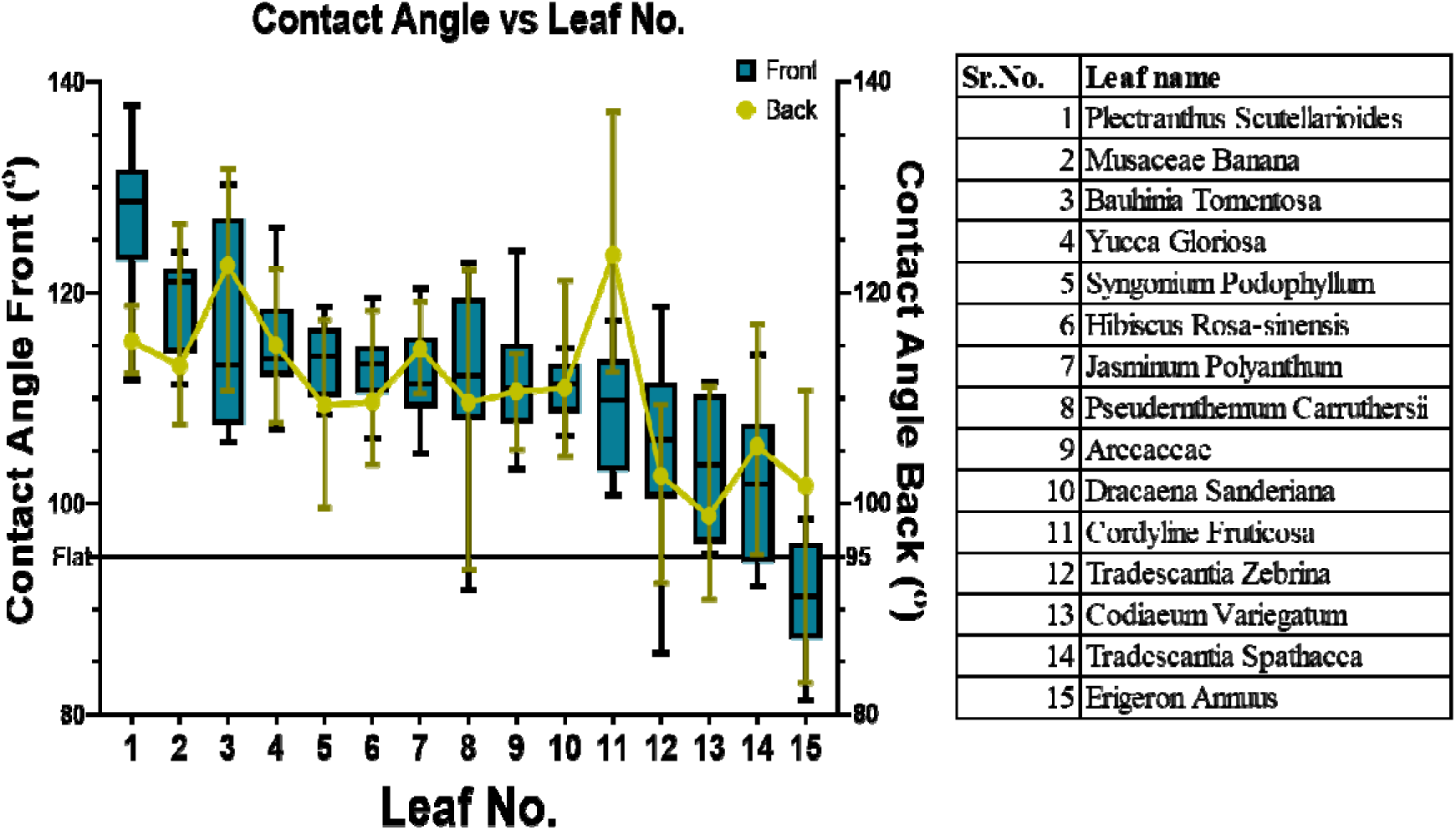
Water contact angle on PDMS substrate for respective leaf template. (as mentioned in table)

Therefore, the morphology and wettability of PDMS changes with respect to the pattern transferred from the leaf template. The structures reported in this work vary widely in terms of feature shape and size. On comparing with wettability, we observed that on the front and back surfaces of some of the leaves the contact angle remained the same even though their surface patterns were different for e.g. for the front side (anisotropic continuous) and back side (anisotropic discontinuous) of *Arecaceae* leaf the contact angle was roughly the same ∼110°. Similarly in the case of *Yucca Gloriosa* leaf (∼115°), which is higher than that of flat PDMS (∼95°). However, with *Cordyline Fruticosa*, the front and back (with anisotropic discontinuous pattern) leaf showed drastically different water contact angle (108° for the front and 124° for the back) even though both are from the same leaf and a similar arrangement of grooves i.e. anisotropic discontinuous pattern. The same was observed in the case of *Erigeron Annuus*. Therefore, to comment on the correlation between bio-mimicked structural properties and wettability, many structures/plant surfaces need to be investigated. Importantly, these variants are produced utilizing readily accessible plant surfaces and straightforward, one-step soft lithography.

For applications requiring high hydrophobicity, we can use the front side of the leaf of *Plectranthus Scutellarioides* (∼127°). Such high hydrophobicity arises from the presence of dense surface hairs, which function as effective water-repellent structures by minimizing the solid–liquid contact area and inhibiting direct interaction between the water droplet and the underlying surface (Otten & Herminghaus, 2004). Based on the same idea the back side of *Bauhinia Tomentosa* leaves have a contact angle of 123°. The anisotropic continuous groove, the back side of *Cordyline Fruticosa*, has a contact angle of 124°. Almost all the leaf surfaces investigated exhibited higher contact angles than flat PDMS (95°), indicating enhanced hydrophobicity. The only exception was the front surface of *Erigeron annuus* (∼91.15°), which is characterized by a reticulate venation pattern and a dense covering of trichomes.

### 3.4. Cell Morphology

The Calcein-AM stained images were analyzed using the NIH ImageJ software to check the alignment of the C2C12 cells cultured on the substrate. Many of the leaves examined possess dense hair coverage, particularly those exhibiting reticulate vein pattern. During the PDMS imprinting process, these hairs tend to fracture and become embedded within the PDMS template. Consequently, leaves with hairy surfaces were excluded from cell study.

For the cell culture study, five surfaces exhibiting the highest water contact angles were selected from leaves with parallel veins, encompassing both anisotropic continuous and anisotropic discontinuous patterns. This selection was based on the higher likelihood of cell alignment with the underlying substrate topography on anisotropic surfaces compared to isotropic patterns or leaves with reticulate vein pattern.

Figure 8 presents anisotropic continuous leaf substrates arranged in order of decreasing contact angle. As shown in the cell projected area and aspect ratio plots (Figure 8g), no clear correlation is observed between the hydrophobicity of the patterned substrates and these cellular properties. Due to the parallel alignment of veins and the presence of rectangular grooves, the anisotropic continuous groove surfaces induce enhanced elongation of C2C12 cells. As a result, C2C12 cells cultured on these substrates exhibit a higher aspect ratio compared to those cultured on flat PDMS (control). Consequently, the absence of topographical constraints on flat PDMS allows cells to spread more freely, resulting in a higher projected cell area compared to cells cultured on parallel, anisotropic continuous grooved surfaces. Out of the five anisotropic continuous pattern surfaces, the front side of *Musaceae Banana* (fig.8 a.i) and the back side of *Dracaena Sanderiana* (fig. 8 d.i) exhibited the highest aspect ratio.

**Fig 8.**
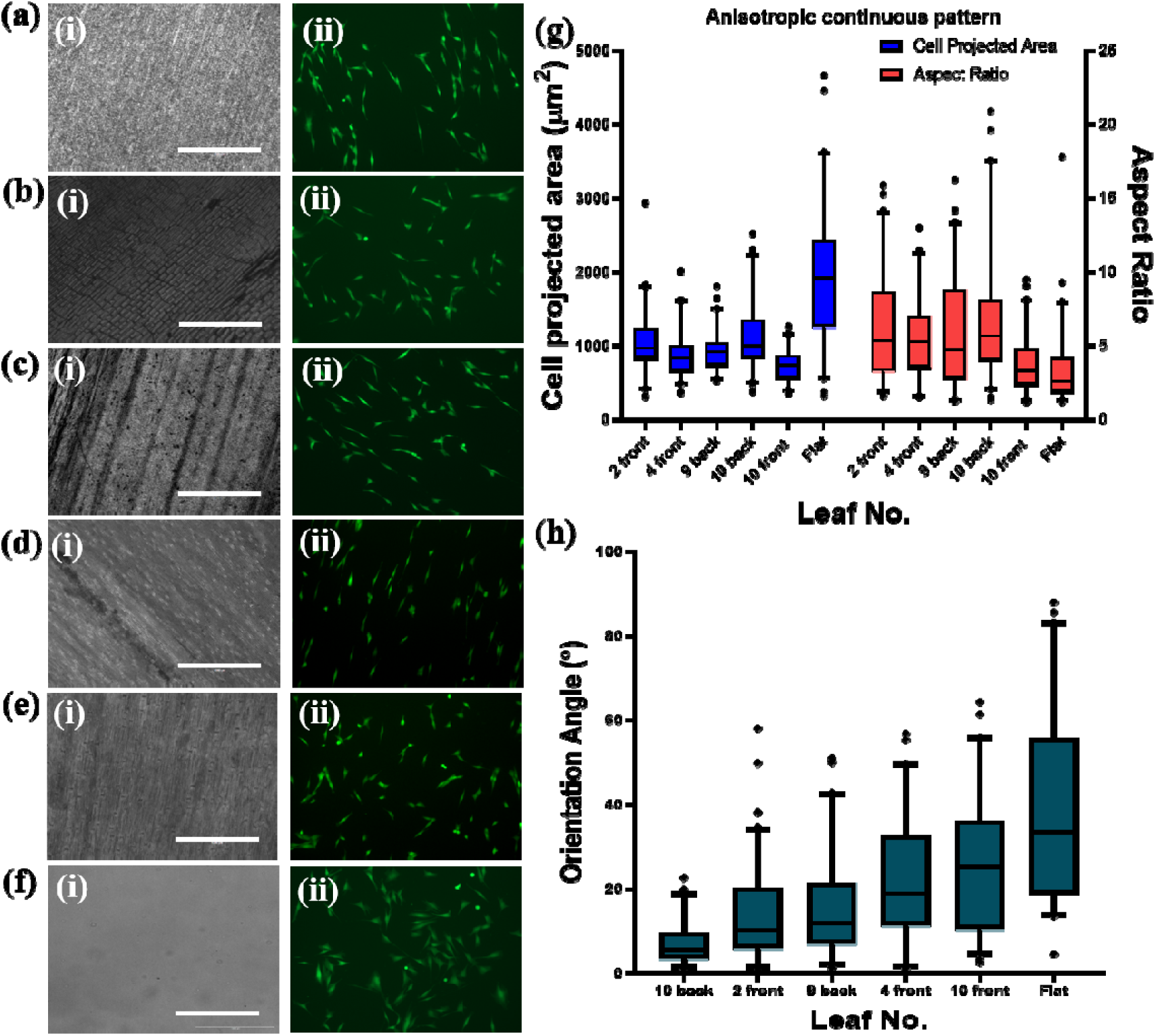
Column 1: Anisotropic continuous pattern microscope image of leaf (Scale is 1000 m) (a) leaf 2 front (b) leaf 4 front (c) leaf 9 back (d) leaf 10 back (e) leaf 10 front (f) Flat PDMS (Refer to Table 1.1 for leaf name) and Column 2: Calcein stained images of C2C12 cells cultured (g) Area and Aspect Ratio of cells cultured on the leaves (h) Orientation Angle of C2C12 cells cultured

A comparative analysis of the trends in fig. 8.g-h, reveals an inverse correlation between cellular morphology and orientation i.e. substrates that promote higher aspect ratios and larger projected cell areas consistently exhibit lower orientation angles. A low orientation angle indicates enhanced alignment of C2C12 cells along the underlying substrate topography. Among the surfaces investigated, the negative replica of the back side of *Dracaena sanderiana* (Fig. 8d.i) induced the highest degree of alignment, as evidenced by the lowest mean orientation angle (7.08°). In contrast, the front side of *Dracaena sanderiana* (Fig. 8e.i) exhibited the weakest alignment, with the highest mean orientation angle (24.58°). C2C12 cells cultured on substrates with anisotropic continuous grooves showed a higher degree of alignment than the control “flat” PDMS substrate.

Figure 9 summarizes the aspect ratio and projected cell area of C2C12 cells cultured on anisotropic discontinuous grooved substrates. Similar to the anisotropic continuous grooves, C2C12 cells cultured on anisotropic discontinuous patterns exhibit a higher aspect ratio and greater degree of alignment of C2C12 cells to the underlying substrate topography than those cultured on flat PDMS. In contrast, the projected cell area of C2C12 cells on flat PDMS is greater than that observed on the grooved substrates. C2C12 cells cultured on the back side of *Musaceae Banana* (fig 9.c) has the highest aspect ratio and the highest degree of alignment (smallest orientation angle = 9.95°) amongst the studied leaves with anisotropic discontinuous grooves. Unlike the anisotropic continuous grooves there is no visible correlation between orientation angle and the aspect ratio/cell projected area.

**Fig 9.**
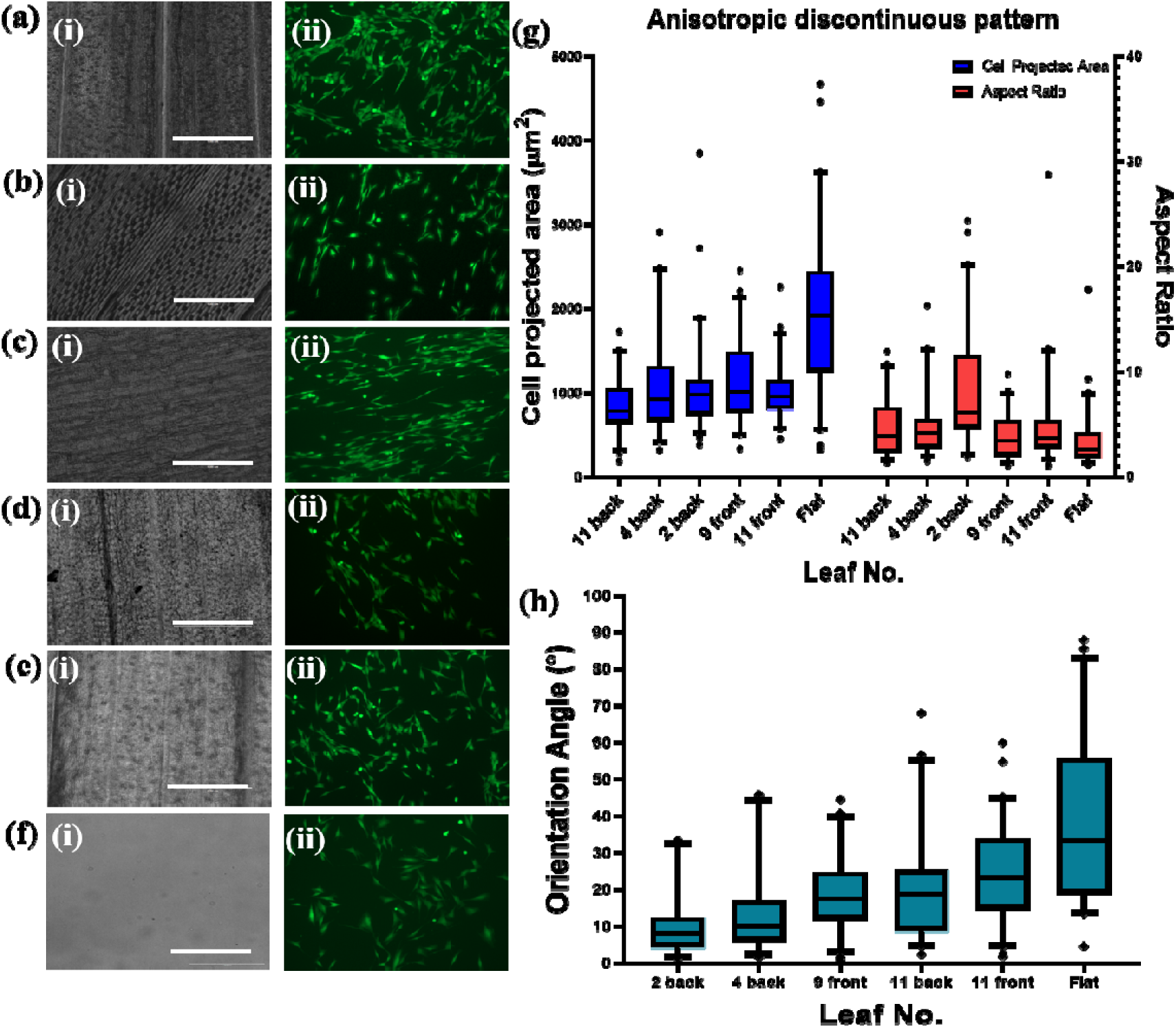
Column 1: Anisotropic discontinuous pattern microscope images of leaves of (Scale: 1000 m) (a) leaf 11 front (b) leaf 4 back (c) leaf 2 back (d) leaf 9 front (e) leaf 11 back (f) Flat PDMS (Refer to Table 1.1 for reference); and Column 2: Calcein stained images of C2C12 cells (g) Area and Aspect Ratio of cells cultured on the leaves (h) Orientation Angle of C2C12 cells cultured

## 4. Conclusion

In this work, we identified and classified the surface topographies of a diverse range of plant leaves and successfully replicated them onto polydimethylsiloxane (PDMS) using a simple soft lithography approach. The replicated surfaces exhibited a wide variety of microstructural features that are difficult to reproduce using conventional micro and nanofabrication techniques, demonstrating the potential of natural templates as an accessible source of complex biomimetic architectures. Notably, many of the plant species investigated in this study have not previously been explored for biomimetic applications, highlighting the vast and largely untapped diversity of natural surfaces that can inspire the design of functional biomaterials.

Among the substrates evaluated for C2C12 cell culture, *Musaceae Banana* had the one of the highest Aspect Ratio in each category, anisotropic continuous and anisotropic discontinuous pattern. Hence, these surfaces serve as a suitable platform for investigating the effect of substrate topography on the differentiation of C2C12 cells into myotubes. From the graphs it is evident that surface hydrophobicity has no clear effect on the aspect ratio or projected cell area of C2C12 cells. However, for the anisotropic continuous pattern, the aspect ratio and projected cell area are positively correlated; as the projected cell area increases, the aspect ratio also increases. Along with this, the orientation angle is inversely related to the aspect ratio/projected cell area for the anisotropic continuous grooves. C2C12 cells cultured on *Dracaena Sanderiana* had the highest (back side) and the lowest (front side) aspect ratio and cell projected area among the anisotropic continuous leaves studied. Both the surfaces also had the highest (back side) and the lowest (front side) degree of alignment among both the category of leaves with parallel veins.

Overall, this work demonstrates that naturally occurring plant surfaces provide a diverse and underexplored library of bioinspired topographies with significant potential for biomaterials research. The proposed classification of leaf-derived surface architectures, supported by FFT analysis, offers a practical framework for comparing complex natural topographies and selecting appropriate templates for specific biomedical applications. The catalogue of replicated plant surfaces presented here can serve as a valuable resource for designing biomimetic substrates that more closely mimic the native cellular microenvironment, with potential applications in organ-on-chip platforms, tissue engineering, and in vitro disease models, ultimately contributing to the development of more physiologically relevant systems for drug screening and regenerative medicine.

